# Hippocampal 4-Hz oscillations emerge during stationary running in a wheel and are resistant to medial septum inactivation

**DOI:** 10.1101/2022.06.20.496892

**Authors:** Ivan Alisson Cavalcante Nunes de Lima, Hindiael Belchior

**Affiliations:** Bioinformatics Multidisciplinary Environment (BioME), Federal University of Rio Grande do Norte, Natal, RN, 59078-970, Brazil; Department of Physical Education, Federal University of Rio Grande do Norte, Natal, RN, 59078-970, Brazil

**Keywords:** 4-Hz oscillations, hippocampus, medial septum inactivation, locomotion

## Abstract

Recent studies described 2-4 Hz oscillations in the hippocampus of rats performing stationary locomotion on treadmills and other apparatus. Since the 2-4 Hz rhythm shares common features with hippocampal theta (5-12 Hz) oscillations - such as a positive running speed-amplitude relationship and modulation of spiking activity - many have questioned whether these rhythms are related or independently generated. To answer this question, we analyzed local field potentials (LFP) and spiking activity from the dorsal CA1 of rats executing a spatial alternation task and running for ∼15 s in a wheel during the intertrial intervals both before and after muscimol injection into the medial septum. We observed 4-Hz oscillations during wheel runs, but not during maze runs. The 4-Hz amplitude positively correlated with wheel speed and negatively with maze speed. Also, the amplitude of 4-Hz and theta oscillations were inversely related. Medial septum inactivation abolished hippocampal theta but preserved 4-Hz oscillations. It also affected the entrainment of pyramidal cells and interneurons by 4-Hz rhythmic activity. In all, these results dissociate the underlying mechanism of 4-Hz and theta oscillations in the rat hippocampus.

**Highlights:**

- 4-Hz oscillations emerge in the rat hippocampus during wheel running and correlate with speed
- 4-Hz and theta amplitude are inversely correlated
- 4-Hz oscillations are resistant to medial septum inactivation
- Rhythmic discharge of interneurons and pyramidal cells at 4-Hz are reduced by medial septum inactivation

**In brief:** Lima and Belchior tested whether the pharmacological inactivation of the medial septum affects hippocampal 4-Hz oscillations associated with stationary running. They found that septal inactivation preserved 4-Hz oscillations while abolished theta, which dissociate the underlying mechanisms of these rhythms in the rat hippocampus.

## INTRODUCTION

The rodent hippocampus expresses abundant rhythmic activity across distinct and sometimes overlapping frequency ranges (Buzsáki and Draguhn, 2004). Theta (5-12 Hz) oscillations are the most prominent rhythm in the rat hippocampus. It rises whenever rats engage in locomotor activity or in response to sensory stimulus (Buzsáki, 2002; Colgin, 2016, 2020). Theta amplitude positively correlates with running speed in mazes, open fields, and other apparatus like stationary running on treadmill and running wheel or passive ambulation (Czurkó et al., 1999; Terrazas et al., 2005; Molter et al., 2012; Li et al., 2014; Cei et al., 2014; Chi et al., 2016; Drieu et al., 2018; Furtunato et al., 2020; Safaryan and Mehta, 2021). Pharmacological experiments established at least two types of hippocampal theta oscillations: an atropine-resistant theta (6-12 Hz, type I) present during locomotor behaviors, and an atropine-sensitive theta (5-9 Hz, type II) expressed in response to sensory stimulation (Vanderwolf, 1969; Kramis et al., 1975). In addition, lesion or inactivation of the medial septum-diagonal band of Broca (MS) abolishes hippocampal theta and impairs memory performance (Green and Arduini, 1954; Stumpf et al., 1962).

In contrast, oscillations at lower frequencies, as in delta (1-4 Hz) band emerge during quiet behaviors and slow-wave sleep episodes, but are typically suppressed during locomotor activity (Buzsáki, 2002; Buzsáki, 2015). The difference between associated behaviors and brain states expressing delta and theta oscillations gave rise to the idea that these two rhythms are essentially orthogonal (Schultheiss et al., 2019). However, recent studies have shown that sustained oscillatory activity in the 2-4 Hz also emerges during locomotion in stationary conditions, like treadmills, head-fixed and virtual reality apparatus (Chi et al., 2016; Furtunato et al., 2020; Safaryan and Mehta, 2021). Hippocampal 2-4 Hz oscillations present similarities with the concomitant theta rhythm, such as a positive relationship between its instantaneous power and running speed, and its phase-modulation of spiking activity (Furtunato et al., 2020; Safaryan and Mehta, 2021). Due to these resemblances, many researchers have raised concerns about the interdependence between the concurrent 2-4 Hz and theta oscillations, and suggested that they may not be clearly dissociated in two genuinely independent rhythms. To sort apart 2-4 Hz oscillations from the classical theta activity and untangle their underlying mechanisms, we analyzed the effects of muscimol injection into the MS over hippocampal oscillations during maze and wheel runs.

## METHODS

The dataset used in this study was previously acquired at the Pastalkova Lab on the Janelia Research Campus (Wang et al., 2017) and made available at http://datadryad.org/ under public domain dedication license.

### Behavioral and electrophysiological recordings

Two 64-channel linear silicon probes (Neuronexus or Janelia RC) were bilaterally implanted at the dorsal CA1 area (coordinates: -4.0 mm AP, ± 3 mm ML) of the rat hippocampus (n = 3 animals across ten sessions). Local field potentials (LFP), spiking activity, and digital video recordings were obtained during a delayed spatial alternation memory task in a U-shaped maze coupled to a running wheel in which animals ran (∼15 s) during the intertrial intervals. Electrophysiological and behavioral recordings were obtained before (n = 304 trials) and after (n = 501 trials) muscimol microinjections into the MS. Detailed experimental procedures can be found in previous publications (Wang et al., 2015; 2017).

### Data analysis

All data analyses were performed using custom made and built-in routines in MATLAB (MathWorks, Natick, MA). First, LFP recordings were notch-filtered between 55 Hz and 65 Hz to remove 60 Hz electrical noise. LFP signals were visually inspected to detect time intervals presenting electrical or movement artifacts. Next, time intervals in which the amplitude of LFP signals were larger than two times the standard deviation were excluded from further analysis (only recordings from rat 943 presented artifacts). Epochs presenting locomotion speed on the maze and in the wheel higher than 10 cm/s were further analyzed.

We used the “eegfilt” function (EEGLAB Toolbox, Delorme and Makeig, 2004) to obtain 4-Hz (3-5 Hz) and theta (6-10 Hz) components of the LFP signals. We refer to 4-Hz oscillations, the spectral frequency between 3-5 Hz. The “pwelch” function from the Signal Processing Toolbox was used to obtain the power spectral density of LFP signals and spiking activity (1-s window length with 90% overlap,Figure 1D). The “hilbert” function from the Signal Processing Toolbox was used to obtain the instantaneous amplitude, phase, and frequency of 4-Hz and theta band components. The “xcorr” function (0.5-s window length, option type “coeff”, Figures 1C and 3B) from the Signal Processing Toolbox was used to obtain the autocorrelograms of LFP signals (Figure 1C and 3B). The “spectrogram” function (2-s window length with 90% overlap, Figure 1B) from the Signal Processing Toolbox to obtain the time-frequency decomposition.

**Figure 1.**
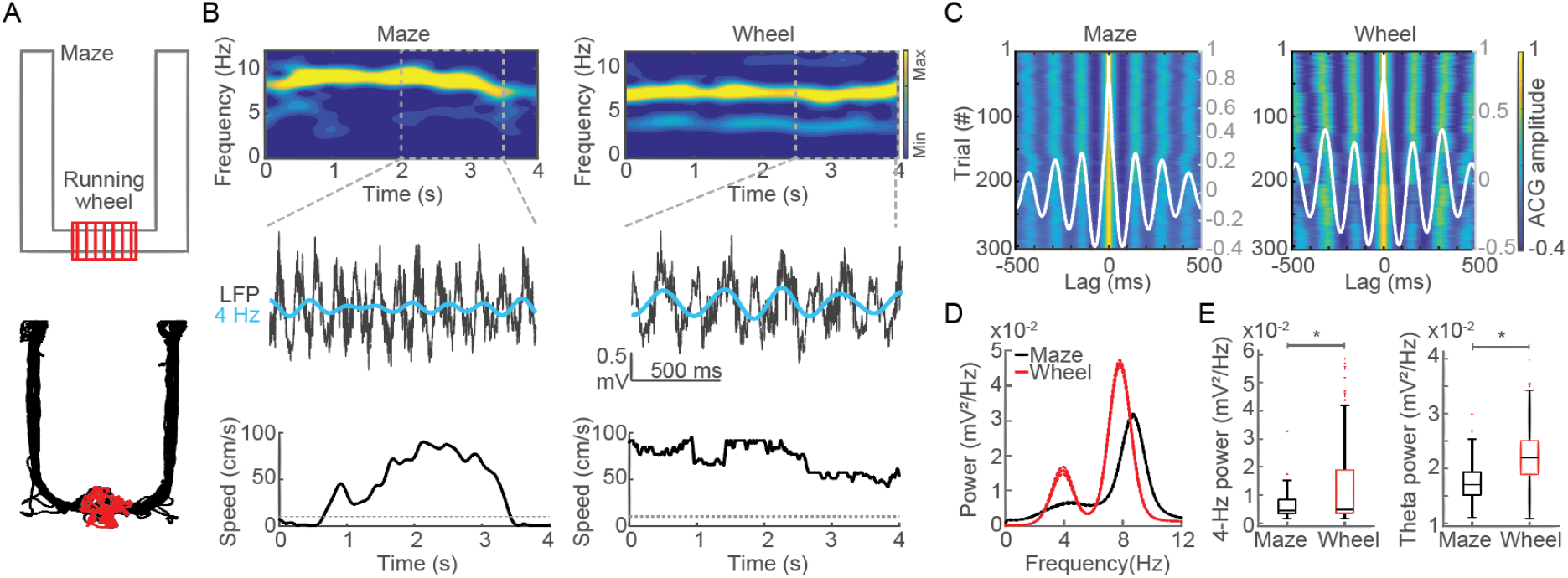
Hippocampal 4-Hz oscillations emerge during wheel but not maze running. (A) Schematic representation of the U-shaped maze (gray) coupled to a running wheel (red, upper). A typical example of the spatial trajectory of a rat on the maze (black) and in the wheel (red, lower). (B) Spectrograms showing energy at 0-12 Hz frequencies during representative maze and wheel runs (left and right, respectively, upper). Notice that only wheel running exhibits prominent energy at 4 Hz and 8 Hz frequencies. Raw LFP and 3-5 Hz band-filtered signals (gray and cyan, respectively, middle), and the instantaneous running speed at the same maze and wheel runs as above (lower). (C) Autocorrelograms of LFP signals across 304 runs at the maze and the wheel (left and right, respectively). Gray traces represent the average autocorrelograms across trials. Interpeak intervals were 145 ms (6.8 Hz) during maze runs and 320 ms (3.1 Hz) during wheel runs. (D) Average power spectra from LFP obtained during maze and wheel runs (black and red, respectively, n=304 trials). Solid lines depict mean and dashed lines depict ± SEM. (E) Boxplot showing the distribution of power at the 3-5 Hz band during maze and wheel runs (left, p < 0.01, WSR test). Distribution of power at the theta (6-10 Hz) band during maze and wheel runs (right, p < 0.01, WSR test).

The rhythmicity of interneurons’ and pyramidal cells’ spiking activity was estimated through autocorrelograms (ACG) and power spectral densities (PSD) of spike times. Only neurons with average firing rate larger than 1 Hz during maze and wheel runs were further analyzed. We used the “xcorr” function (0.5-s window length, option type “coeff”, Figure 4A) to calculate the ACG of each neuron individually. The ACG of each neuron was normalized before obtaining the group result: NormACG = [ACG - min(ACG)] / [max(ACG) - min(ACG)], where max(ACG) and min(ACG) denote the maximum and minimum ACG values of each neuron, respectively. Only neurons with normalized ACG larger than a threshold value of 0.2 were further analyzed. To obtain the group result, the normalized ACGs were then averaged by cell type and condition (Figure 4A). To graphically display the ACG of each neuron (Figure 4A), we used a color coded plot of the z-scored data smoothed by 100 points, in which warm colors mean higher ACG values for a given neuron at a given time lag. The values of ACG peak amplitudes and interpeak intervals of each neuron were obtained within specific time windows for 4-Hz (lags at 200 - 300 ms) and theta (lags at 100 - 200 ms) bands.

Similarly, we used the “pwelch” function (5-s time window, and no overlap, Figure 4B) to estimate the PSD of each neuron individually. Only the neurons previously analyzed in the ACG were included in PSD analyses. The relative PSD of each neuron was obtained before compute the group result: relative PSD = [PSD / sum(PSD)], in which sum(PSD) denotes the sum of power across frequency values. The group result shown in Figure 4B was obtained by averaging the relative PSD by cell type and condition. Similarly to previous studies (Wang et al., 2015), the spike PSD peaks were at higher frequencies than previously observed to LFP PSD (Figure 1D). We thus adjusted the 4-Hz and theta frequency bands to fit the oscillatory spiking activity for 4-6 Hz and 8-15 Hz, respectively. Since different conditions exhibited different levels of baseline power, we calculated the power index to compare across conditions. For instance, the 4-Hz power index = max(PSD value at 4-6 Hz band) - PSD value at 6 Hz, in which max(PSD) denotes the maximum power value within the band of interest.

### Statistical analysis

All statistical analyses were performed in MATLAB. An alpha level of 0.05 was used to denote statistical significance. Group data are expressed as mean ± standard error of the mean (SEM) or median and quartiles over trials. We used the Shapiro-Wilk test to evaluate data normality. The Wilcoxon signed-rank (WSR) test or paired t-test was used to compare between maze and wheel running conditions. The Wilcoxon rank-sum (WRS) test or Student’s t-test was used to compare between wheel runs pre and post muscimol injections, and between wheel runs previous to correct and incorrect choices. The “corr” function (option type “Spearman”) from the Signal Processing Toolbox was used to obtain the Spearman’s rank-based correlation coefficients (rho) between speed-amplitude and speed-frequency at the 4-Hz and theta bands (5-s time window, Figures 2B and 2D), and also between the instantaneous amplitude of coexisting 4-Hz and theta oscillations (5-s time window, Figures 2C and 2E).

**Figure 2.**
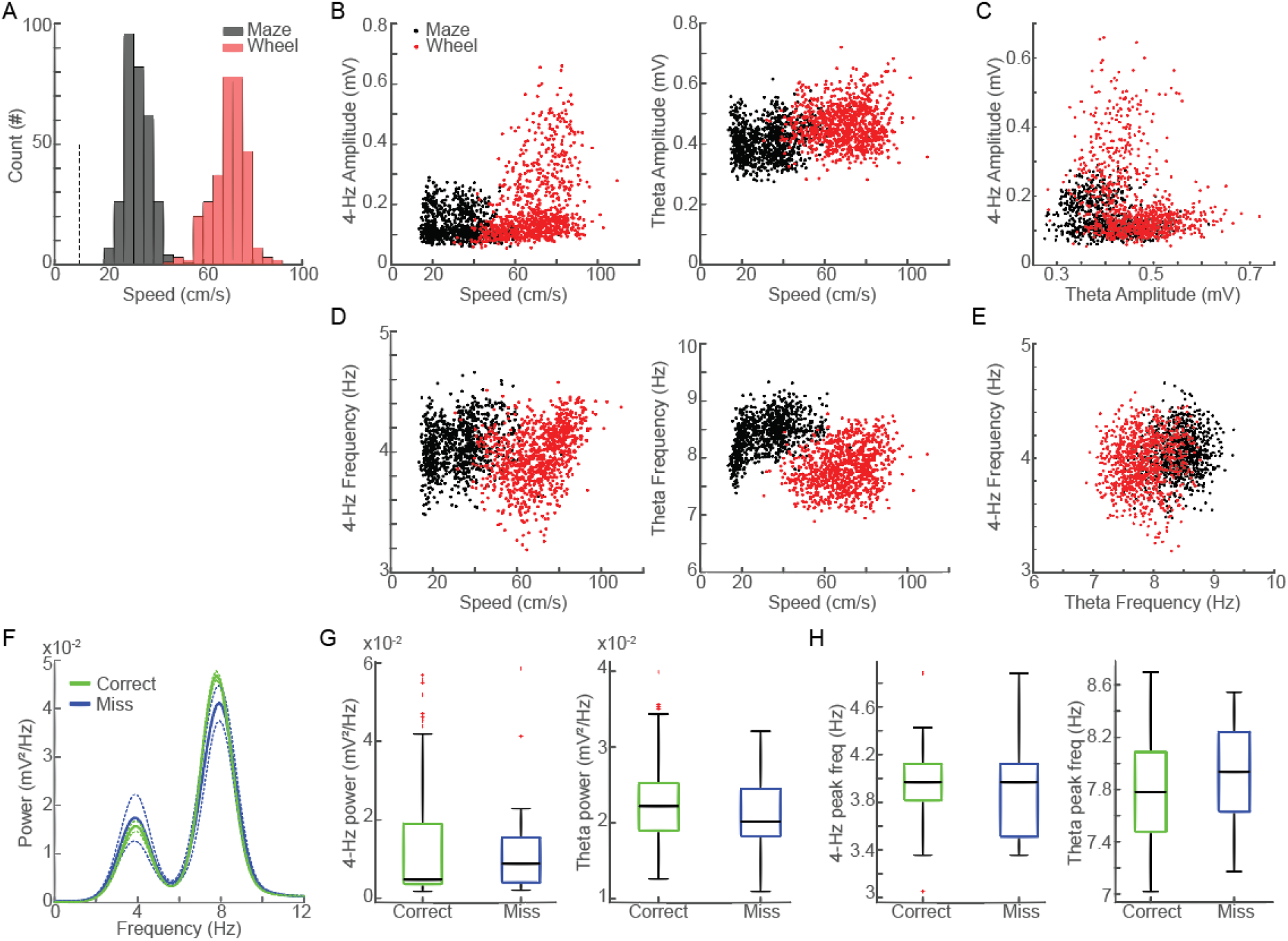
Hippocampal 4-Hz oscillations are modulated by running speed but are not affected by choice performance. (A) Histograms of running speeds at the maze (black) and at the wheel (red, n = 304 trials, p < 0.01, WRS test). (B) Scatter plots of running speed and the instantaneous 4-Hz amplitude (left, wheel: rho = 0.38, p < 0.01, maze: rho = -0.15, p < 0.01) and theta amplitude (right, wheel: rho = 0.15, maze: rho = 0.01, p = 0.74, p < 0.01). Notice the inverse relationship between 4-Hz amplitudes at maze and wheel conditions. (C) Relationship between 4-Hz and theta amplitude on the maze (black, rho = -0.31, p < 0.01) and in the wheel (red, rho = -0.28, p < 0.01). (D) Scatter plots show the relationship between running speed and the instantaneous 4-Hz frequency (left, 4-Hz at the wheel: rho = 0.38, p < 0.01; 4-Hz at the maze: rho = 0.27, p < 0.01) and theta frequency (right, theta at the maze: rho = 0.46, p < 0.01; theta at the wheel: rho = 0.23, p < 0.01). (E) Relationships between the 4-Hz and theta instantaneous peak frequency on the maze (black, rho = 0.11, p < 0.01) and in the wheel (red, rho = 0.10, p < 0.01). (F) Average power spectra during wheel run previous to correct (green, n = 286 trials) and incorrect (blue, n = 18 trials) choices at the spatial alternation memory task. Solid lines represent means and dashed lines represent ± SEM. (G) Boxplots of 4-Hz band power (left, p = 0.26, WRS test) and theta band power (right, p = 0.34, WRS test) during wheel runs previous to correct and incorrect choices. (H) Boxplots of 4-Hz peak frequency (left, p = 0.75, WRS test) and theta peak frequency (right, p = 0.36, WRS test) during wheel runs before correct and incorrect trials (see also histograms at Figure 2 - figure Supplement 1).

## RESULTS

### 4-Hz oscillations emerge in the rat hippocampus during wheel running

Rats performed a spatial alternation task on a U-shaped maze and ran for ∼15 s on a wheel during the intertrial intervals (Figure 1A). Consistent with previous reports, spectral decomposition of CA1 LFP showed prominent theta oscillations while rats ran on both maze and wheel (Figure 1B). Interestingly, however, only wheel runs further exhibited remarkable rhythmicity at 4-Hz. Hippocampal 4-Hz oscillations were noticeable at the raw LFP and spectrograms during wheel but maze runs, even in trials with similar running speeds (Figure 1B and Figure 1 - figure Supplement 1). Autocorrelograms revealed strong rhythmicity during maze and wheel runs with mean interpeak intervals of 145 ms (6.8 Hz) and 320 ms (3.1 Hz), respectively (Figure 1C). Average power spectra at the group level showed a single peak at 8.7 Hz during maze runs and two peaks at 7.8 Hz and at 4 Hz in the wheel (Figure 1D). The 4-Hz power was significantly higher during wheel than maze runs (Figure 1E, left), and its peak frequency was lower at the wheel (Figure 1 - figure Supplement 2A). The theta power was higher at the wheel (Figure 1E, right), while its peak frequency was slower at the wheel (Figure 1 - figure Supplement 2B).

### Hippocampal 4-Hz amplitude correlates with running speed in the wheel

Animals ran faster at the wheel (Figure 2A), which could suggest that 4-Hz oscillations are due to higher running speeds. In fact, the instantaneous 4-Hz amplitude and running speed were positively correlated at the wheel (Figure 2B, left), and negatively correlated at the maze. In turn, the theta amplitude was positively correlated with maze speed (Figure 2B, right) but not with wheel speed. The instantaneous amplitude of 4-Hz and theta oscillations were negatively correlated (Figure 2C). Since 4-Hz oscillations were only observed during the intertrial intervals of the spatial alternation task, we next evaluated whether 4-Hz power and frequency in the wheel were associated with memory performance. Wheel runs previous to correct and incorrect choices exhibited similar power spectra (Figure 2F), in which neither 4-Hz nor theta band power nor peak frequency were statistically different (Figure 2G).

### Hippocampal 4-Hz oscillations are resistant to medial septum inactivation

Next, we evaluated how MS inactivation affects 4-Hz and theta oscillations in the wheel. Muscimol injection reduced running speed on wheel (Figure 3A), with no changes in the duration of wheel runs (post: 13.43 s, p = 0.06, WRS test). It also impaired choice performance at the spatial alternation task (Figure 3 - figure Supplement 1). Autocorrelograms and power spectral analysis show that muscimol injection affected theta rhythmicity but surprisingly preserved 4-Hz oscillations (Figure 3B and 3C). Muscimol injections did not change 4-Hz power and significantly decreased theta power (Figure 3D). In addition, muscimol injections reduced both 4-Hz and theta peak frequency (Figure 3E).

**Figure 3.**
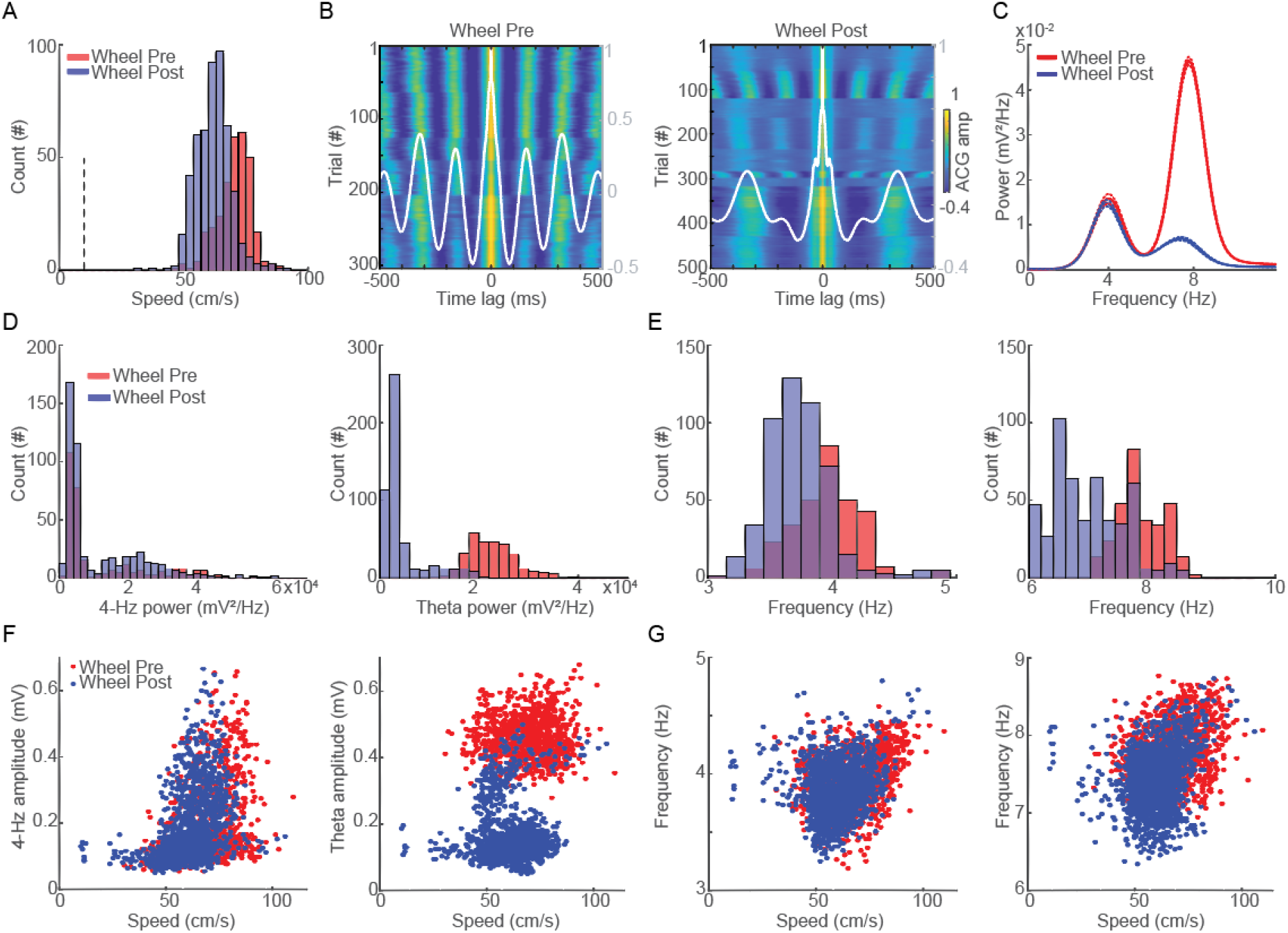
Hippocampal 4-Hz oscillations are resistant to pharmacological inactivation of the medial septum. (A) Histograms of running speed at the wheel before (red, n = 304 trials, 13.54 s) and after (blue, n = 501 trials, 13.43 s) muscimol inactivation of the MS (p < 0.01, WRS test). (B) Color-coded autocorrelograms of LFP signals across runs at the wheel before (left) and after (right) muscimol. Gray traces represent the average autocorrelograms. (C) Average power spectra at 0-12 Hz during wheel runs before (red) and after (blue) muscimol injection. Solid lines represent the mean and dashed lines represent ± SEM. (D) Histograms of 4-Hz band power (left, p = 0.68, WRS test) and theta band power (right, p < 0.01, WRS test) before and after muscimol. (E) Histograms of 4-Hz band peak frequency (left, p < 0.01, WRS test) and theta band peak frequency (right, p < 0.01, WRS test) before and after muscimol. (F) Scatter plots of running speed and the instantaneous amplitude of 4-Hz (left, pre: rho = 0.38, p < 0.01; and post: rho = 0.52, p < 0.01) and theta (right, pre: rho = 0.01, p = 0.74; and post: rho = 0.08, p < 0.01) oscillations before and after muscimol. (G) Scatter plots of running speed and the instantaneous frequency of 4-Hz (left, pre: rho = 0.38, p < 0.01; and post: rho = 0.19, p < 0.01) and theta (right, pre: rho = 0.23, p < 0.01; and post: rho = 0.29, p < 0.01) oscillations before and after muscimol.

We then evaluated whether the effects of MS inactivation on 4-Hz and theta oscillations were dependent on running speed in the wheel. Muscimol injection did not affect the 4-Hz amplitude-running speed relationship (Figure 3F, left). In contrast, muscimol injection reduced theta amplitude even at identical running speeds (Figure 3F, right). Moreover, muscimol injection did not affect peak frequency neither at 4-Hz (Figure 3G, left) nor at theta oscillations (Figure 3G, right). Muscimol abolished the relationship between 4-Hz and theta instantaneous amplitudes in the wheel (Figure 3 - figure Supplement 2A). Muscimol also affected the relationship between 4-Hz and theta instantaneous peak frequency in the wheel (Figure 3 - figure Supplement 2B).

### 4-Hz oscillations modulate spiking activity of interneurons and pyramidal cells

We evaluated autocorrelograms and power spectra of spiking activity of putative interneurons and pyramidal cells during maze and wheel runs. Corroborating previous studies, autocorrelograms of spiking activity during maze runs presented strong rhythmicity at the theta band (Figure 4A, upper panel). During wheel runs, however, spike ACG exhibited remarkable rhythmicity at the 4-Hz band frequency (Figure 4A, middle panels). In this condition, interneurons presented longer interpeak intervals and higher amplitudes at 4-Hz than in maze runs. Pyramidal cells also showed longer interpeak intervals and higher ACG amplitudes at the 4-Hz band frequency during wheel than maze runs (Figure 4 - figure Supplement 1). At the 4-Hz band frequency, muscimol injection did not significantly affect neither interneurons’ ACG amplitude nor interpeak intervals. Muscimol reduced the ACG amplitude of pyramidal cells, but did not change interpeak intervals in the 4-Hz frequency.

**Figure 4.**
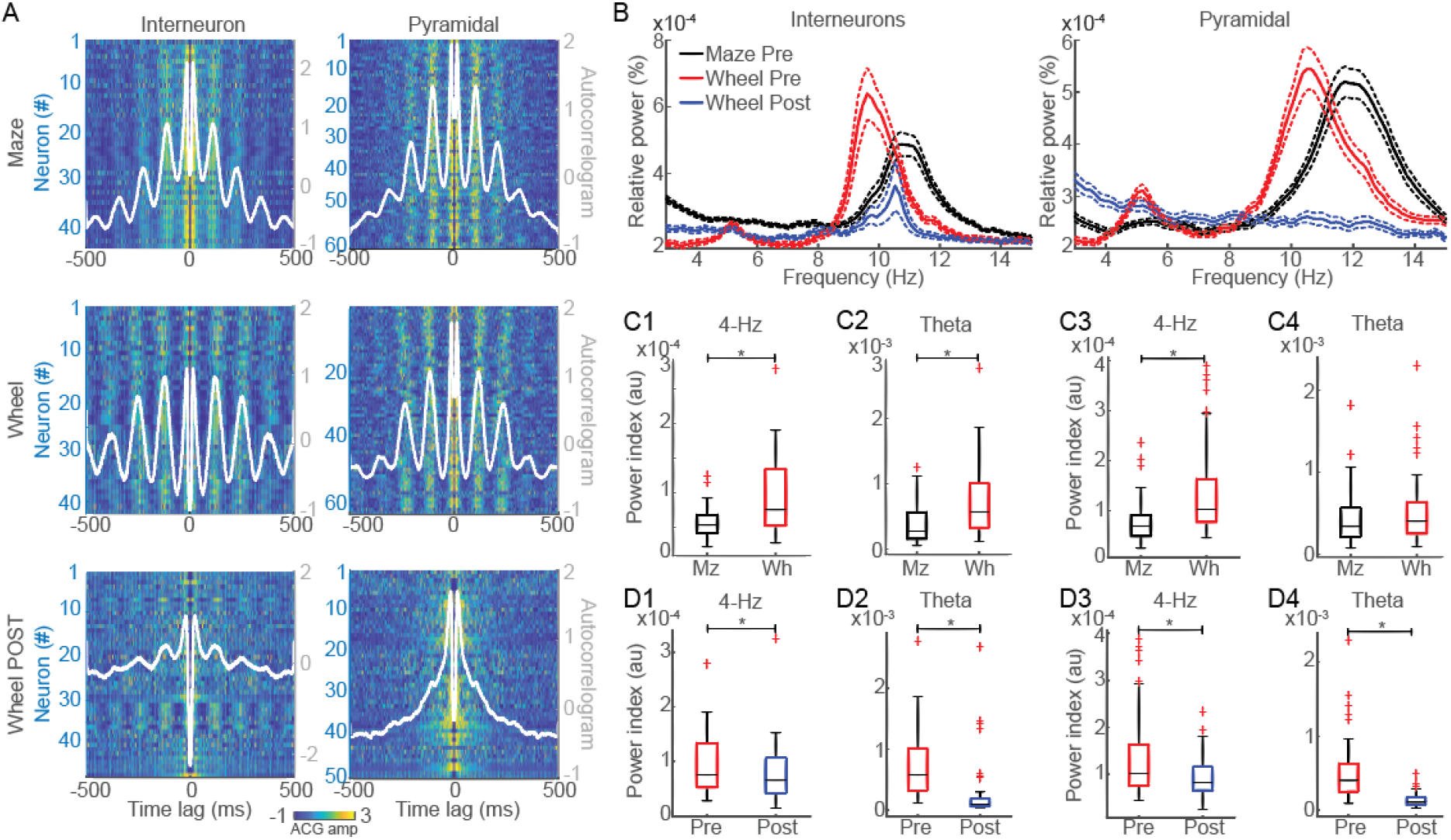
4-Hz rhythmicity of spiking activity decreases after MS inactivation. (A) Color-coded autocorrelograms of interneurons (left) and pyramidal cells (right) during maze runs (upper), and during wheel runs before (middle) and after muscimol injection (lower). White lines depict the average ACG for the neuronal population. Notice that neuronal rhythmicity of both cell types increased during wheel runs and strongly decreased after muscimol injection. Interneurons interpeak intervals at 4-Hz in the wheel: 246 ms; on the maze: 227 ms (p < 0.01, WRS test). Interneurons ACG amplitude at 4-Hz in the wheel: 0.94; on the maze: 0.73 (p < 0.01, WRS test). Pyramidal cells interpeak intervals at 4-Hz in the wheel: 232 ms; on the maze: 218 ms (p < 0.01, WRS test). Pyramidal cells ACG amplitudes in the wheel: 0.66; on the maze: 0.5 (p < 0.01, WRS test). Interneurons interpeak intervals at 4-Hz in the wheel before: 0.94; after muscimol: 0.9 (p = 0.42, WRS test). Interneurons ACG amplitudes at 4-Hz in the wheel before: 246 ms; after muscimol: 250 ms (p = 0.37, WRS test). Pyramidal cells interpeak intervals at 4-Hz in the wheel before: 232 ms; after muscimol: 232 ms (p = 0.94, WRS test). Pyramidal cells ACG amplitudes at 4-Hz in the wheel before: 0.66; after muscimol: 0.5 (p < 0.01, WRS test). (B) Relative (%) power spectral density of interneurons’ (left) and pyramidal cells’ (right) spiking activity during maze runs (black), wheel runs before (red), and wheel runs after muscimol injection (blue). (C) Boxplots comparing spike PSDs of interneurons (left) and pyramidal cells (right) on 4-Hz and theta bands during maze and wheel runs. (D) Same as in panel C for wheel runs before (pre) and after (post) muscimol injection. * p < 0.05, and ** p < 0.01 at WRS or WSR tests.

Power spectra of interneurons’ and pyramidal cells’ spikes during maze runs exhibited a single peak at the theta band frequency. In contrast, wheel runs exhibited two peaks: one at the theta band and a second around 4-Hz (Figure 4B). Interneurons and pyramidal cells showed higher power index at 4-Hz in wheel than maze runs (Figure 4C1 and C3, respectively). In contrast, only interneurons presented higher power index at theta in wheel than maze runs, not pyramidal cells (Figure 4C2 and C4). Finally, muscimol injection significantly reduced interneurons’ and pyramidal cells’ power index in the 4-Hz (Figure 4 D1 and D3) and in the theta (Figure 4 D2 and D4).

## DISCUSSION

We report the emergence of 4-Hz oscillations in the dorsal CA1 of rats engaged in stationary running in a wheel. Hippocampal 4-Hz oscillations directly correlate with wheel speed and entrain rhythmic spiking activity of pyramidal cells and interneurons. Differently from theta rhythms, 4-Hz oscillations are resistant to pharmacological inactivation of the medial septum. Our results disentangle the mechanisms of generation of 4-Hz and theta oscillations in the rat hippocampus.

Crescent evidence has shown that during stationary locomotion the rat hippocampus can simultaneously exhibit two distinct rhythms within the 1-12 Hz frequency range: the classical 5-12 Hz theta and a new 2-5 Hz oscillation specifically observed during locomotion on treadmills, virtual reality and head-fixed apparatus (Chi et al., 2016; Furtunato et al., 2020; Safaryan and Mehta, 2021). Our findings confirm the concurrency of hippocampal 4-Hz and theta oscillations during stationary runs, extending to wheel apparatus.

Interestingly, only 4-Hz amplitude displayed a positive relationship with running speed in wheel (rho = 0.31, theta rho = 0.01, respectively). Their instantaneous amplitudes were inversely related (rho = -0.28), which differed from Safaryan and Mehta (2021) in a virtual reality apparatus. These results are at least partially consistent with Furtunato et al. (2020) that described an inverse relationship between the power of theta and 2-4 Hz oscillations, in which theta power decreased and 2-4 Hz power increased across consecutive runs at same speed (30 cm/s) on a treadmill. Despite their concomitant occurrence, the orthogonality between these rhythms may indicate a potential dissociation in the mechanism of generation of hippocampal theta and 4-Hz oscillations.

To evaluate this possibility, we tested whether pharmacological inactivation of MS through intracerebral microinjections of muscimol affected hippocampal 4-Hz oscillations. As expected (Winson, 1978; Wang et al., 2015), muscimol injections abolished hippocampal theta. However, hippocampal 4-Hz oscillations were resistant to MS inactivation and preserved a positive relationship with running speed in the wheel. These results provide the first evidence of different mechanisms of generation of 4-Hz and theta oscillations occurring in the rat hippocampus during stationary runs.

It is still unclear why 4-Hz oscillations are expressed in stationary locomotion but not usually observed during translational conditions. Previous studies presented preliminary data of speed-modulated 4-Hz during wheel runs (Czurko et al., 1999; Molter et al., 2012). However, none of them directly compared stationary versus translational locomotion. Here, in the same recordings only wheel runs were accompanied by 4-Hz oscillations but not speed-matching maze runs. It confirms findings by Safaryan and Mehta (2021) comparing runs in a linear track and in a virtual reality apparatus. In addition, since our behavioral protocol tested wheel runs during the intertrial intervals of a spatial alternation task, we were capable to evaluate if 4-Hz oscillations in the wheel were associated with choice performance. Neither the power nor the peak frequency at 4-Hz in the wheel differed before correct and incorrect choices, suggesting that 4-Hz oscillations were not associated with cognitive demands of the spatial alternation task.

Theta oscillations dominate the hippocampus during translational locomotion, with amplitude and frequency positively associated with running speed and acceleration (Kropff et al., 2021; Kennedy et al., 2022). When rats engage in stationary running, however, the absence of linear body movements decouples vestibular and proprioceptive signals, which may dissociate hippocampal theta in two components at 4 and 8 Hz (Safaryan and Mehta, 2021). Alternatively, hippocampal oscillations may also synchronize with the respiratory rhythm (Chi et al., 2016; Lockmann et al., 2016). Chi et al. (2016) showed that mice running in head-fixed conditions exhibited long periods of steady respiration around 4-Hz that entrain this rhythm in the dentate gyrus. The respiratory frequency is more stable during stationary than translational locomotion because of the reduction of bouts of sniffing, which could also explain why 4-Hz oscillations are not observed during maze runs. Future research that simultaneously monitors breathing and hippocampal rhythms in stationary and translational runs could investigate this hypothesis.

Finally, the present work provides pharmacological and electrophysiological evidence that hippocampal 4-Hz and theta oscillations are independently generated during stationary locomotion.

## ACKNOWLEDGEMENTS

Supported by Conselho Nacional de Desenvolvimento Científico e Tecnológico (CNPq: 409753/2021) and Coordenação de Aperfeiçoamento de Pessoal de Nível Superior (CAPES). We thank Pastalkova’s laboratory for making data publicly available at http://datadryad.org/.

## DATA AVAILABILITY

The raw electrophysiological and behavioral data analyzed in the current study are publicly available via Dryad (https://doi.org/10.5061/dryad.j021s). Algorithms used throughout this study are publicly available at Github (https://github.com/belchiorlab/Lima-Belchior2022.git).

## AUTHOR CONTRIBUTIONS

H.B. designed research; I.A.C.N.L. and H.B. analyzed data; I.A.C.N.L. and H.B. wrote the manuscript.

## LIST OF FIGURE SUPPLEMENT

**Figure 1 - figure Supplement 1.**
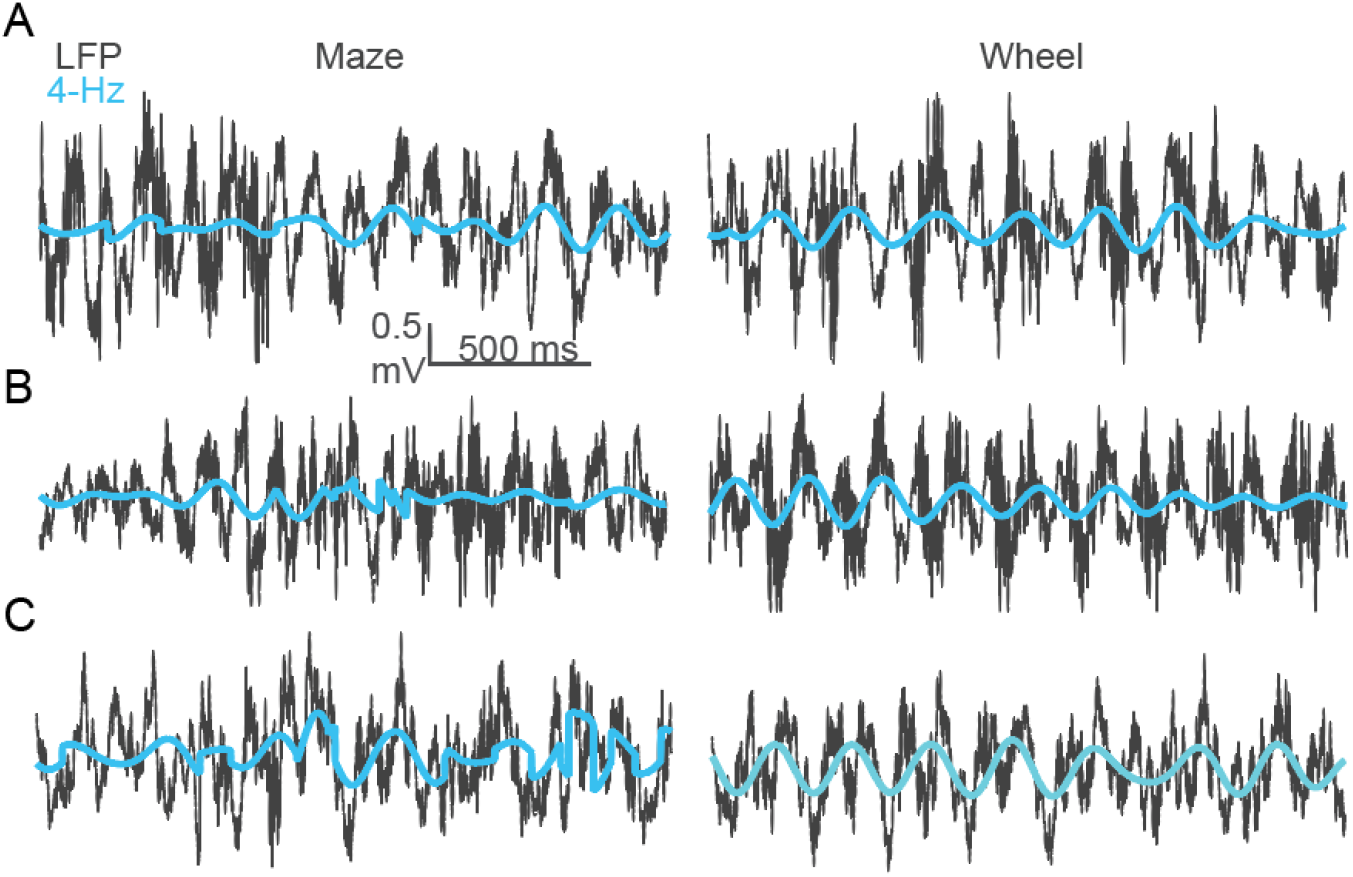
4-Hz oscillations are noticed at the raw LFP signals during wheel runs. Representative examples of raw LFP (black) and 4-Hz-filtered signals (cyan) recorded during maze (left) and wheel runs (right). Panels A, B and C depict signals from three different rats (rat A498-20120807-01, rat A543-20120412-01, and rat A943-20120526-01, respectively). Only epochs of running speed larger than 10 cm/s are shown.

**Figure 1 - figure Supplement 2.**
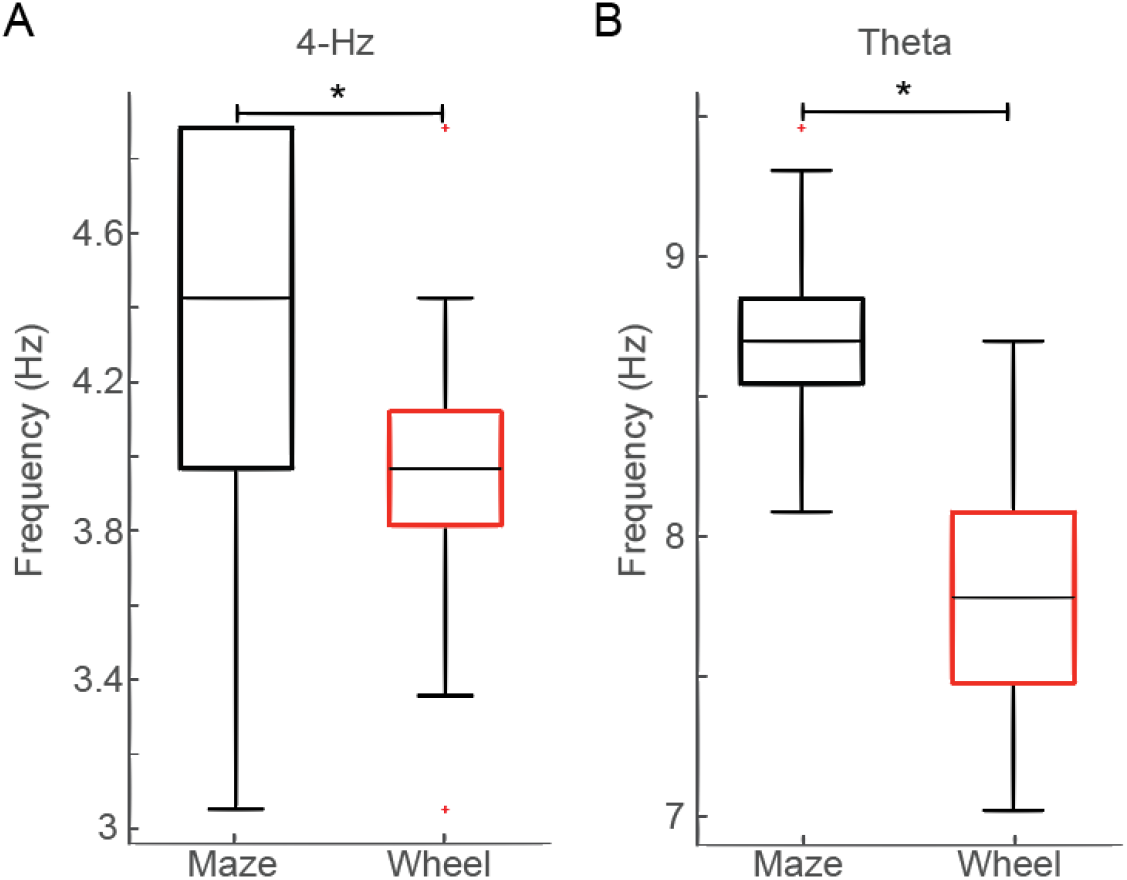
Peak frequency within 4-Hz and theta bands during maze and wheel runs. (A) Peak frequency within the 4-Hz band during maze and wheel runs (p < 0.01, WSR test, n = 304 trials). (B) Peak frequency within the theta band during maze and wheel runs (p < 0.01, WSR test, n = 304 trials).

**Figure 2 - figure Supplement 1.**
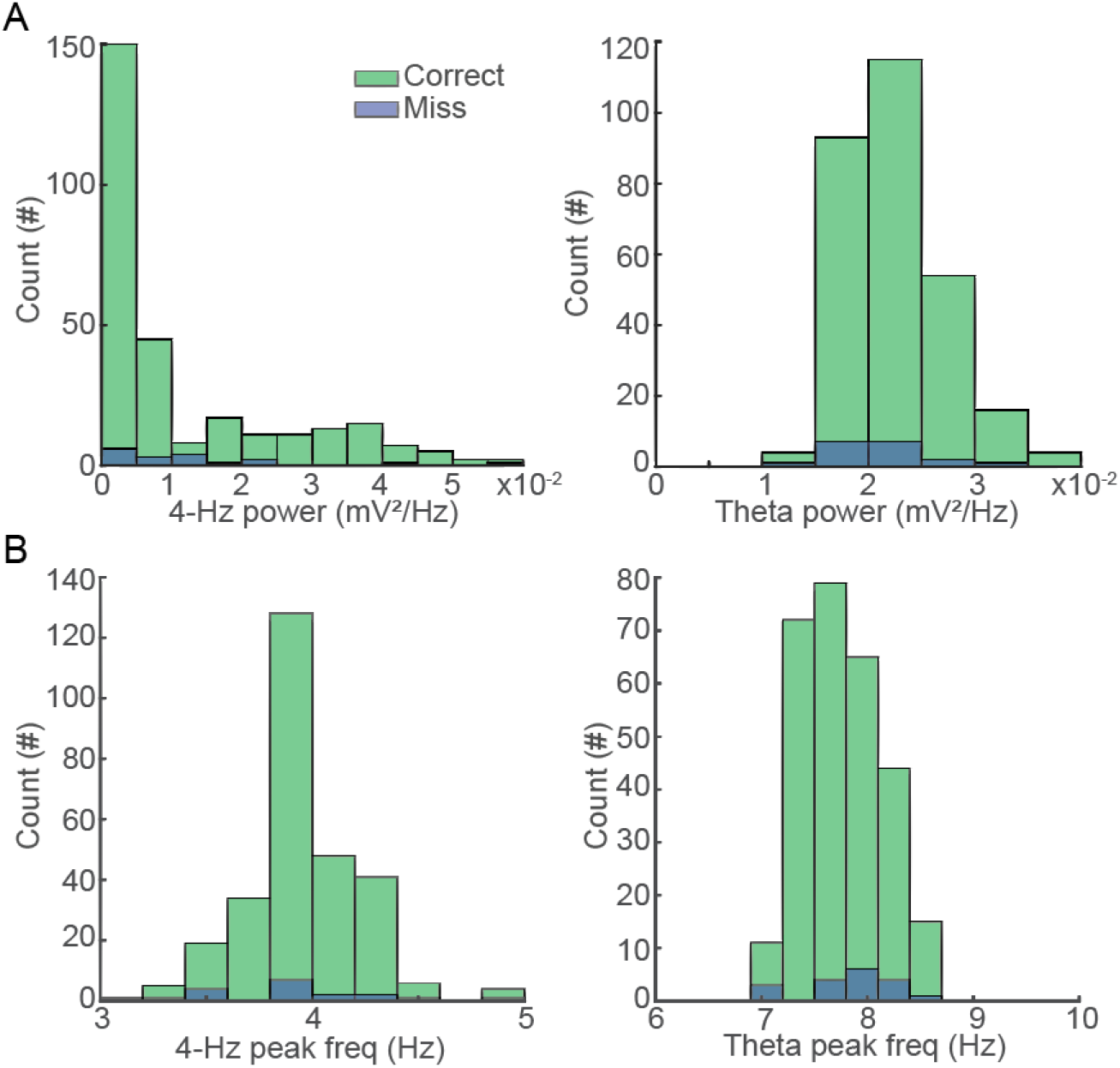
Power and peak frequency at 4-Hz and theta bands during wheel runs according to choice performance on the task. (A) Histograms of 4-Hz band power (left) and theta band power (right) during wheel runs previous to correct (green) and incorrect (blue) choices. (B) Distribution of 4-Hz peak frequency (left) and theta peak frequency (right) during wheel runs before correct (green) and incorrect (blue) trials.

**Figure 3 - figure Supplement 1.**
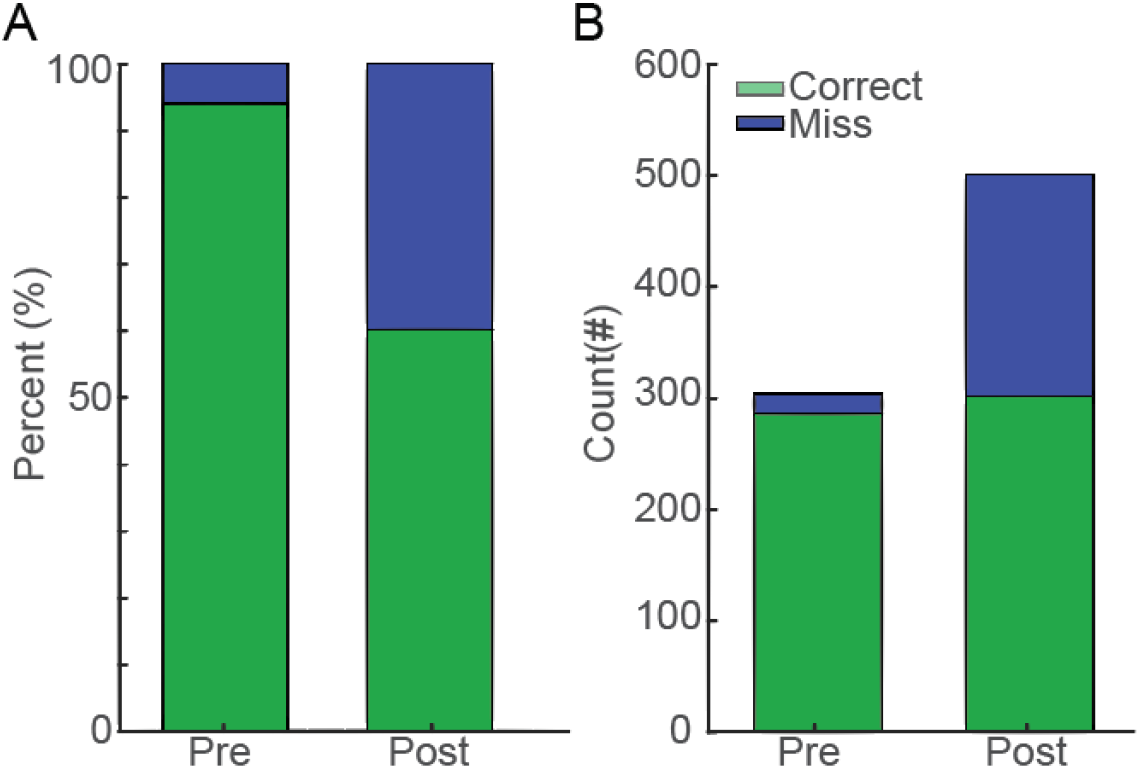
Choice performance before and after muscimol injection in the MS. (A) Percentage of correct choices before (pre, correct 94.07%, green, from 304 trials) and after (post, correct 60.27%, green, from 501 trials) muscimol administration into the MS (across 10 sessions from 3 animals, p < 0.01, WRS test). (B) Absolute number of correct (green) and incorrect (blue) choices before (pre, correct 286 trials) and after (post, correct 302 trials) muscimol injection.

**Figure 3 - figure Supplement 2.**
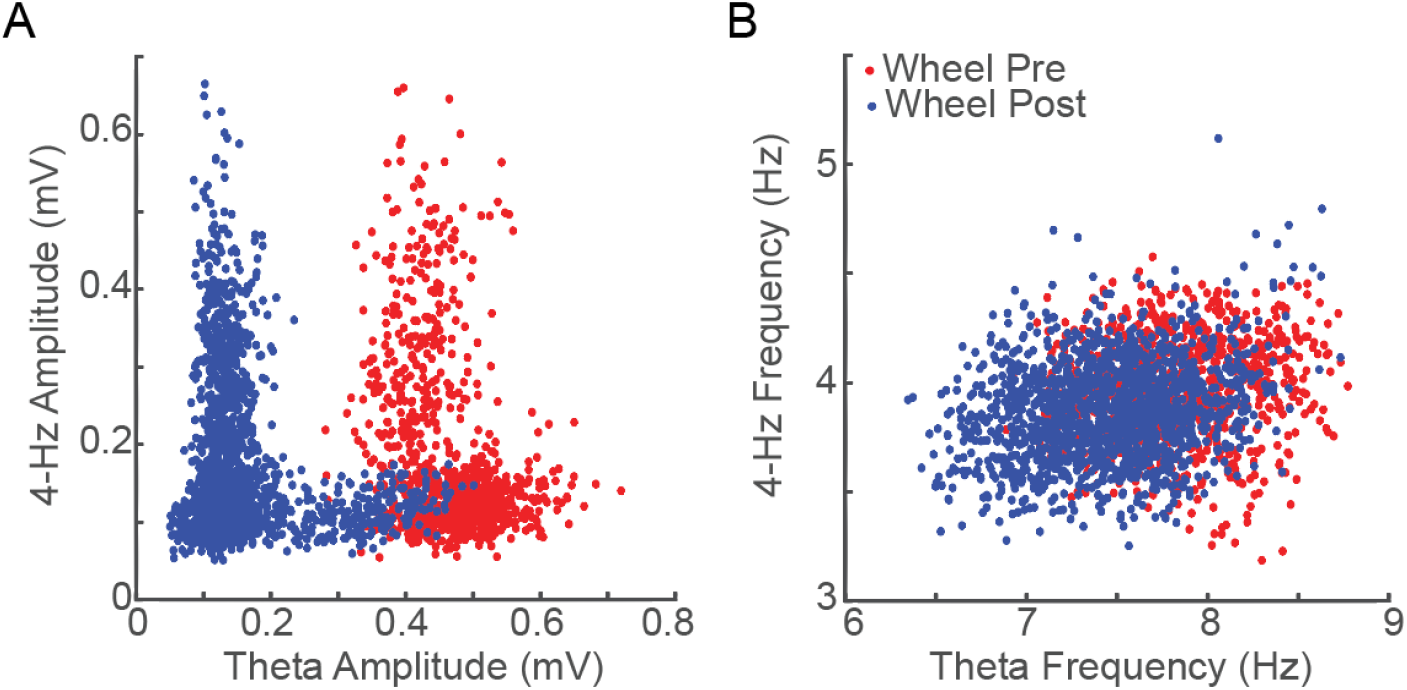
Relationship between 4-Hz and theta instantaneous amplitude and frequency before and after muscimol injection. (A) Relationship between 4-Hz and theta amplitude in the wheel before (pre, red, rho = -0.28, p < 0.01, n = 908 5-s bins) and after (post, blue, rho = -0.04, p = 0.08, n = 1540 5-s bins) muscimol injection. (B) Relationship between 4-Hz and theta peak frequency in the wheel before (pre, red, rho = 0.10, p < 0.01, n = 908 5-s bins) and after (post, blue, rho = 0.19, p < 0.01, n = 1540 5-s bins) muscimol injection.

**Figure 4 - figure Supplement 1.**
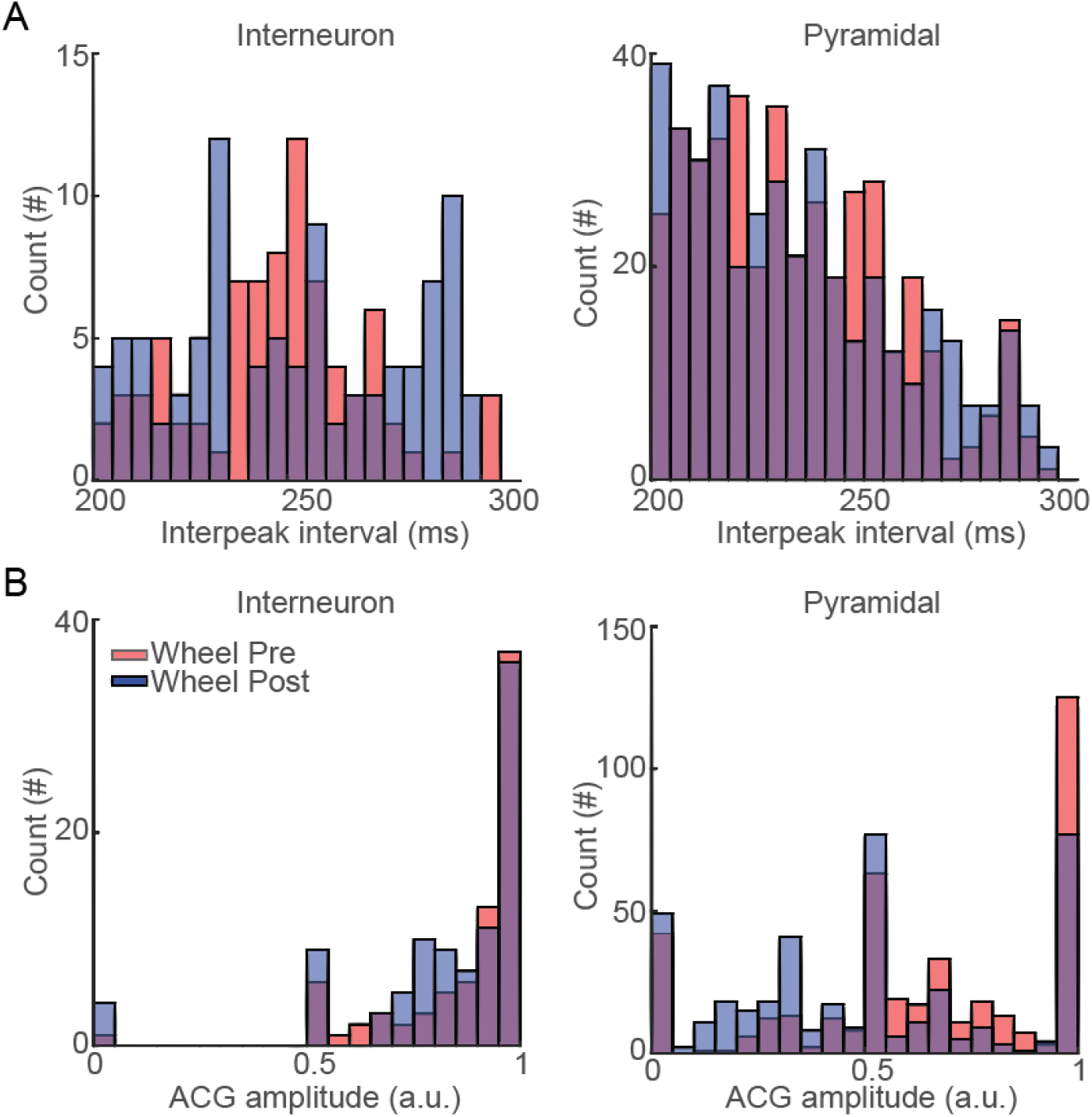
Amplitude and interpeak intervals of spike ACGs at 4-Hz in the wheel before and after muscimol injection. (A) ACG amplitudes of interneurons’ (left) and pyramidal cells’ (right) spikes at the 4-Hz frequency range. (B) ACG interpeak intervals of interneurons’ (left) and pyramidal cells’ (right) spikes at the 4-Hz frequency range. Red and blue bars depict wheel runs before and after muscimol injections, respectively.

